# The taxonomic and functional biogeographies of phytoplankton and zooplankton communities across boreal lakes

**DOI:** 10.1101/373332

**Authors:** Nicolas F. St-Gelais, Richard J. Vogt, Paul A. del Giorgio, Beatrix E. Beisner

## Abstract

Strong trophic interactions link primary producers (phytoplankton) and consumers (zooplankton) in lakes. However, the influence of such interactions on the biogeographical distribution of the taxa and functional traits of planktonic organisms in lakes has never been explicitly tested. To better understand the spatial distribution of these two major aquatic groups, we related composition across boreal lakes (104 for zooplankton and 48 for phytoplankton) in relation to a common suite of environmental and spatial factors. We then directly tested the degree of coupling in their taxonomic and functional distributions across the subset of common lakes. Although phytoplankton composition responded mainly to properties related to water quality while zooplankton composition responded more strongly to lake morphometry, we found significant coupling between their spatial distributions at taxonomic and functional levels based on a Procrustes test. This coupling was not significant after removing the effect of environmental drivers (water quality and morphometry) on the spatial distributions of the two groups. This result suggests that top-down and bottom-up effects (e.g. nutrient concentration and predation) drove trophic interactions at the landscape level. We also found a significant effect of dispersal limitation on the distribution of taxa, which could explain why coupling was stronger for taxa than for traits at the scale of this study, with a turnover of species observed between regions, but no trait turnover. Our results indicate that landscape pelagic food web responses to anthropogenic changes in ecosystem parameters should be driven by a combination of top-down and bottom-factors for taxonomic composition, but with a relative resilience in functional trait composition of lake communities.

## Introduction

The observed composition of ecological communities is the result of multiple assembly processes (Kraft and Ackerly 2014) occurring at different spatial scales (Declerck et al. 2011). Locally, species interactions (e.g. competition, predation, mutualisms) form a dominant assembly mechanism structuring ecological communities (Diamond 1975). However, the effect of local species interaction on the distribution of organisms at large scales, within and across regions, is not well understood (Gotelli et al. 2010, Wiens 2011) - including for plankton communities across lakes. Within a lake, planktonic consumers (zooplankton) and primary producers (phytoplankton) interact strongly, mainly via predation (Porter 1977, Sterner 1989), and studies on trophic cascades have emphasized how changes at one trophic level can affect entire food webs (Carpenter et al. 1985). While the top-down and bottom-up effects in lake food webs (e.g. predation effects of zooplankton but also of fish on zooplankton; nutrient effects on phytoplankton, but also of phytoplankton availability on zooplankton) have typically been studied for their influence on standing biomass, they also influence community composition (McQueen et al. 1989, Ghadouani et al. 2003). However, the degree to which these strong local trophic interactions affect the larger-scale distribution of zooplankton and phytoplankton across lake metacommunities at regional scales is still unknown.

To our knowledge, very few studies have examined the joint distributions of phytoplankton and zooplankton. Such studies would help to reveal the importance of the trophic link between these community constituents as a potential constraining factor of their respective taxonomic distributions on a landscape. Soininen and colleagues (2007) compared the spatial concordance of distance-decay relationships of phytoplankton and zooplankton taxa across a series of ponds, finding that they were significantly stronger for zooplankton than for phytoplankton, but only across drainage basins and not within basins. However, they did not directly assess the joint taxonomic distributions of the different plankton groups.

Although zooplankton exhibit prey selectivity (Vanderploeg 1981, Paffenhöfer 1984), individual taxa often feed on many different phytoplankton species (Knisely and Geller 1986). Zooplankton grazing thus reflects selection for certain functional traits (e.g. size, shape) that are common to a number of phytoplankton species, more than for specific taxonomic groups per se. Thus, if trophic interactions between phytoplankton and zooplankton are strongly constrained by functional traits of both plankton groups, then joint biogeographical concordances should be more evident for functional traits than for strict taxonomic classification. A temporal analogue of this idea is the repeatable seasonal succession observed in many temperate lakes and captured effectively by the Plankton Ecology Group (PEG) model, which characterizes the dynamic effects of such trophic interactions by referring to specific functional groups of phytoplankton and zooplankton (Sommer et al. 1986, 2012). If functional trophic interactions are equally spatially important, and repeatable, across a landscape, one would expect any biogeographical concordance between phytoplankton and zooplankton to be more observable using a functional trait lens than with a taxonomic one. Functional traits currently available for lake plankton account mechanistically for trophic interaction potential related to food web responses, including resource availability (phytoplankton), feeding behaviours (zooplankton), and predator evasion (phytoplankton and zooplankton, Barnett et al. 2007, Hébert et al. 2016).

Unlike trophic interactions, the effect of habitat on the taxonomic biogeography of zooplankton and phytoplankton is well-studied at the regional scale (see Keller and Pitblado 1989, Pinel-Alloul et al. 1990, 1995, O’Brien et al. 2004, Soininen et al. 2011, Stomp et al. 2011). However, the relative importance of drivers appears to differ between the two groups. Phytoplankton respond strongly to water chemistry, and especially to nutrient concentrations (Pinel-Alloul et al. 1990; Duarte et al. 1992; Tolonen et al. 2020). For example, cyanobacteria and diatoms tend to dominate in eutrophic lakes (Watson et al. 1997), which also impacts the functional trait composition of the phytoplankton community. On the other hand, while zooplankton also respond to water chemistry (Pinel-Alloul et al. 1995) the response to lake characteristics, like depth and macrophyte cover, is often stronger (Tolonen et al. 2020), probably reflecting the top-down effect of fish predation on zooplankton (Rodriguez et al. 1993). Hence, based on the concordance between the distribution of phytoplankton and zooplankton and the strength of the response of both groups to top-down (e.g. lake depth) and bottom up factors (e.g. nutrients), it is possible to assess the relative importance of trophic interactions between zooplankton and phytoplankton on their spatial distribution.

Like environmental gradients, dispersal limitation is also known to structure phytoplankton and zooplankton communities to differing degrees (Heino et al. 2015). Previous work has indicated that phytoplankton generally respond to environmental factors while zooplankton are more dispersal-limited (Beisner et al. 2006, De Bie et al. 2012, Padial et al. 2014). Such differential response patterns could also interfere with the emergence of a strong biogeographical coupling of the groups. In fact, differential relative responses of phytoplankton and zooplankton to environmental factors and distances between lakes, make it necessary to control for their respective effects before assessing the importance of trophic interactions on the biogeographical coupling of the two plankton groups.

We conducted the first comprehensive study that directly considers interactions between planktonic organisms at two trophic levels and their joint biogeographical responses to environmental conditions and dispersal (based on distances). Understanding of the importance of trophic interactions at the landscape-scale can guide interpretation of the effects of broad-scale anthropogenic changes on aquatic food webs (e.g. Jeziorski et al. 2015). To this end, we assessed the strength of the coupling between the taxonomic and functional distribution of lake zooplankton and phytoplankton communities, given that spatially coherent trophic interactions between the groups would result in a consistent co-occurrence of species and functional traits, To deepen our understanding of the underlying drivers of spatial coherence, we used a stepwise framework in which we assessed the extent to which the observed coupling could be explained by environmental gradients and then by limitation to dispersal. If any observed coupling is explained by variation in the environment (water quality or morphometry) it would suggest that, in a way analogous to seasonal succession (temporal change) in lakes, local food web interactions in response to environmental gradients structure the distribution of both groups together (bottom-up or top-down interactions). However, if any observed coupling is explained by space instead of environment, it would suggest that similar rates of dispersal limitation, instead of local interactions, are the main driver of coupling between the two groups. Finally, if any coupling observed is not explained by environment or space, it would suggest that coupling results mostly from pure trophic interactions that are not structured by environmental gradients. This stepwise approach also allowed us to investigate the environmental and spatial factors influencing each group (phytoplankton or zooplankton) separately as well.

Our study covers a large biogeographical range (the maximum distance between any two lakes is 1 228 km) and examines more than 100 lakes clustered in three regions that together characterize geological and environmental variation in the boreal belt across one of Canada’s largest provinces (Québec). Because trophic interactions are mediated by functional traits, we expected a stronger residual coupling between the distribution of zooplankton and phytoplankton traits compared to any taxonomic coupling. Furthermore, we expected coupling to be even stronger when only traits specific to trophic interactions between zooplankton and phytoplankton were considered (e.g. pigment type, feeding strategy).

## Methods

### Study lakes and sampling

Crustacean zooplankton samples were collected from 104 lakes with a low level of anthropogenic disturbance, within three environmentally (Figure 1, Table 1) and geologically (Roy 2012) distinct regions of Quebec province, Canada, during the years 2010 (Abitibi, May to October), 2011 (Chicoutimi, June to October) and 2012 (Schefferville, July to August). Integrated zooplankton samples were collected from the deepest point of each lake using a conical plankton net (110 µm mesh, 0.30 m mouth diameter), equipped with a flow meter (General Oceanics, USA), hauled vertically from 1 m above the sediments to the surface. Zooplankton samples were anaesthetized using carbonated water and were preserved in 75% (final concentration) ethanol. In a subset of 48 lakes, the phytoplankton community was also simultaneously sampled over the photic zone using a flexible PVC sampler tube and an integrated subsample (250 ml) was preserved in Lugol’s solution.

**Figure 1:**
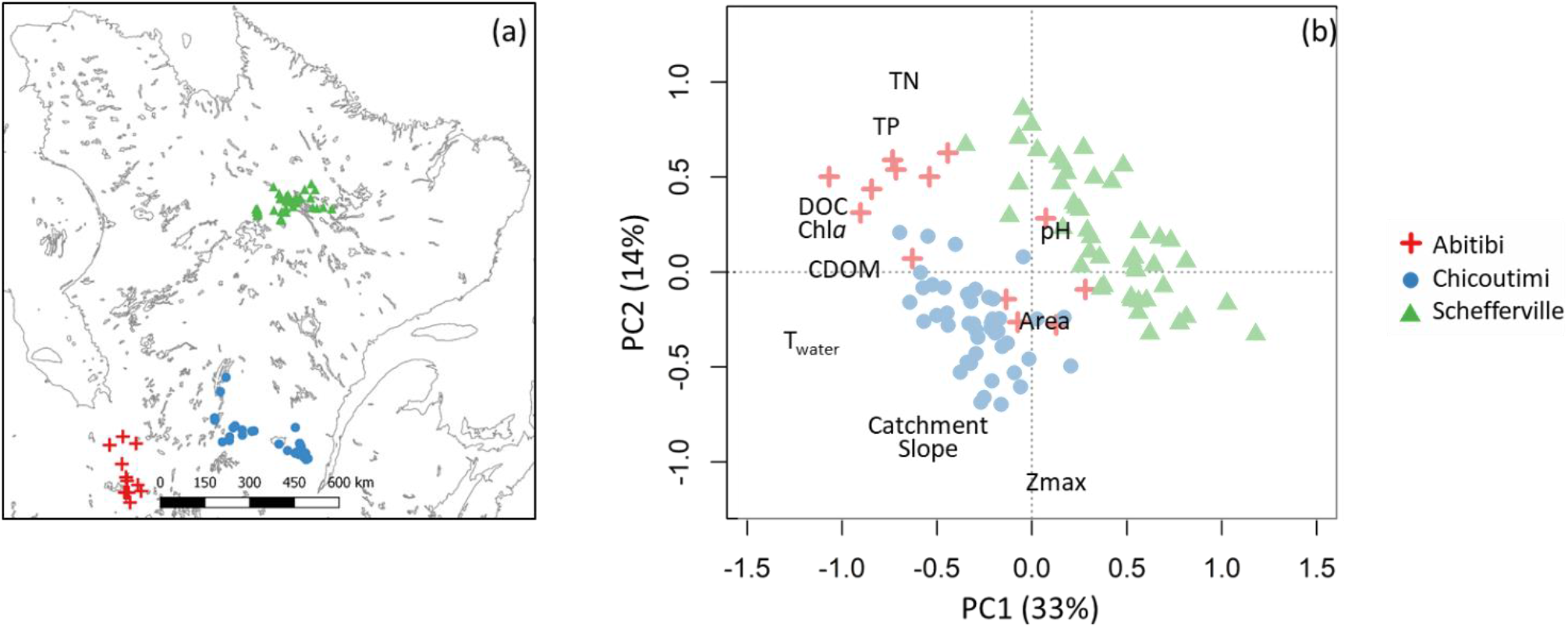
(a) Map of sampled lakes across three regions of boreal Quebec, Canada. (b) Lakes environmental principal component analysis (PCA) based on water quality and morphometric variables where the different symbols represent the three sampled regions. Abbreviations are as follows total phosphorus (TP); total nitrogen (TN); dissolved organic carbon (DOC); chlorophyll a (*Chl-a*); coloured dissolved organic matter (CDOM); water temperature (Twater); lake maximum depth (Zmax); lake area (Area).

**Table 1:** Means and standard error (SE, in parenthesis) of environmental variables: water quality (WQ), and morphometry (M) with the number of missing values (#NA) and the transformation used.

| Variable name<br>(abbreviation) | Category | #<br>NA | Units | Transformation | Regional averages<br>(Standard Error) |  |  |  |  |  |
| --- | --- | --- | --- | --- | --- | --- | --- | --- | --- | --- |
|  |  |  |  |  | Abitibi |  | Chicoutimi |  | Schefferville |  |
|  |  |  |  |  | Phyto. | Zoo. | Phyto. | Zoo. | Phyto. | Zoo. |
| Chlorophyll a<br>(Chl-a) | WQ | 0 | ug/L | log<sub>10</sub> | 3.1<br>(0.7) | 3.7<br>(0.9) | 1.9<br>(0.2) | 2.0<br>(0.1) | 0.9<br>(0.1) | 0.8<br>(0.1) |
| Total Phosphorus<br>(TP) | WQ | 0 | ug/L | log<sub>10</sub> | 17.0<br>(3) | 18.5<br>(3.5) | 9.8<br>(0.9) | 10.1<br>(0.6) | 7.3<br>(0.7) | 7.7<br>(0.6) |
| Total Nitrogen<br>(TN) | WQ | 0 | mg/L | log<sub>10</sub> | 0.3<br>(0.05) | 0.3<br>(0.04) | 0.2<br>(0.02) | 0.2<br>(0.0<br>1) | 0.2<br>(0.02) | 0.2<br>(0.01) |
| Dissolved organic carbon<br>(DOC) | WQ | 2 | mg/L | log<sub>10</sub> | 12.4<br>(1.4) | 12.4<br>(1.3) | 7.5<br>(0.7) | 7.7<br>(0.4) | 3.8<br>(0.4) | 4.0<br>(0.2) |
| Colored dissolved organic matter (CDOM) | WQ | 1 | 1/m | $\log_{10}$ | 3.2<br>(0.7) | 3.4<br>(0.8) | 3.9<br>(0.5) | 4.0<br>(0.4) | 1.0<br>(0.2) | 1.1<br>(0.1) |
| pH | WQ | 1 | - | - | 7.4<br>(0.3) | 7.5<br>(0.3) | 6.9<br>(0.1) | 7.0<br>(0.1) | 6.8<br>(0.2) | 7.1<br>(0.1) |
| Water temperature (0.5 m) ($T_{\text{water}}$) | WQ | 1 | $^{\circ}\text{C}$ | - | 21.8<br>(1.1) | 20.6<br>(1.0) | 20.0<br>(0.5) | 19.8<br>(0.3) | 14.2<br>(0.2) | 15.0<br>(0.2) |
| Maximum depth (Zmax) | M | 0 | m | $\log_{10}$ | 5.7<br>(1.4) | 5.7<br>(1.2) | 8.2<br>(1.5) | 9.4<br>(1.1) | 7.3<br>(1.5) | 6.8<br>(0.9) |
| Lake Area (Area) | M | 0 | $\text{m}^2$ | $\log_{10}$ | 5.89<br>(0.3) | 5.7<br>(0.3) | 6.1<br>(0.3) | 6.2<br>(0.2) | 6.0<br>(0.3) | 6.1<br>(0.2) |
| Number of lakes |  |  |  |  | 11 | 13 | 16 | 44 | 21 | 46 |

Crustacean zooplankton were identified at the species level (but aggregated for analyses at the genus level to correspond to the phytoplankton data), using an inverted microscope (50-400X) and individuals were counted until a total of 200 individuals had been enumerated. For each taxon present in a lake, the length of 20 mature individuals was measured and biomass by taxon was estimated using length-dry-mass regressions (McCauley 1984, Culver et al. 1985). Phytoplankton were enumerated at the genus level using the Ütermohl method on an inverted microscope at 400X magnification. Phytoplankton biomass was estimated from biovolume computed using cell and colony length measurements and corresponding geometric forms (Hillebrand et al. 1999). We also measured key limnological variables to characterize the lake and catchment environments. We used a multiparameter sonde (YSI, Yellow Springs Instruments, OH, USA) to measure pH (at 0.5m depth) and temperature (at 0.5m depth intervals, then averaged over the water column). Water samples were collected at 0.5m depth to measure the concentration of chlorophyll a (Chl-a), total phosphorus (TP), total nitrogen (TN), dissolved organic carbon (DOC) and coloured dissolved organic matter (CDOM). Chl-a was extracted with 90% hot ethanol and absorption was measured spectrophotometrically before and after acidification to account for phaeophytin (Lorenzen 1967, Nush 1980); TP was measured from water samples using the molybdenum-blue method following persulfate digestion (Cattaneo and Prairie 1995); TN was measured using nitrates after persulfate digestion; DOC concentration was measured on an O.I. Analytical (Texas, USA) TIC/TOC using 0.45 μm filtered water after sodium persulfate digestion; CDOM was measured using a UV/Vis UltroSpec 2100 spectrophotometer (Biochrom, Cambridge, UK) at 440 nm. Missing values (see Table 1) were imputed using an approach based on random forest (missForest R package, Stekhoven and Bühlmann 2012, NRMSE : 0.038). Lake depth was measured at sampling point using a Portable Water Depth Sounder Gauge (Cole-Parmer). Lake area was derived using ArcGIS V10 software (ESRI Inc., Redland, CA, USA) and catchment slope was estimated using a Digital Elevation Model (Canadian Digital Elevation Data). To visualize environmental differences between the three regions, we used a principal component analysis (PCA) using the rda function (vegan R package, Oksanen et al. 2015). Finally, because our sampling was discontinuous on the landscape we used the Euclidean distance between lakes to characterize the effect of dispersal limitation on the distribution of taxa and functional traits.

### Functional trait composition and diversity

Given that our objective was to test for a significant coupling between adjacent trophic groups, we selected functional traits (Table 2) that explicitly characterize the grazing interaction between phytoplankton and zooplankton, as well as their interaction with other trophic levels in the food web. For phytoplankton, we selected traits that define how they interface with resources (i.e. nutrients and light) and the zooplankton grazers (motility, edibility, colony formation). For zooplankton, we focused on traits that define how they consume phytoplankton (i.e. feeding type and trophic group). Phytoplankton trait values were obtained from a literature review (see (Longhi and Beisner 2010 for details) and included (i) capacity for N-fixation, (ii) silica demand, (iii) capacity for mixotrophy, (iv) pigment composition (v) cell motility and (vi) edibility to zooplankton (>35 μm linear dimension) and (vii) tendency to form colonies. Crustacean zooplankton traits values were also obtained from a literature review (see Barnett et al. 2007) and included (i) feeding type and (ii) trophic group. For both zooplankton and phytoplankton, we also used the average individual biomass of each taxon as an integrative functional trait of body size (Brown et al. 2004, Litchman et al. 2013) related both to resource acquisition and grazer avoidance (Table 2).

**Table 2:** Phytoplankton and zooplankton functional traits used in this study.

|  | Category | Traits | Values |
| --- | --- | --- | --- |
| Phytoplankton | Resource acquisition | Nitrogen fixation | yes/no |
|  |  | Si requirement | yes/no |
|  |  | Mixotroph | yes/no |
|  |  | Pigments | Green<br>Blue<br>Brown<br>Mixed |
|  | Grazer avoidance | Motility | None<br>F(Flagellated)<br>V(Vacuole) |
|  |  | Biovolume | numerical |
|  |  | Colonial | yes/no |
| Zooplankton | Resource acquisition | Trophic Position | Carnivore<br>Omnivore-Carnivore<br>Omnivore<br>Omnivore-Herbivores<br>Herbivores<br>Immature |
|  |  | Feeding Type | B (Bosmina)-filtration<br>C (Chydorus)-filtration<br>D (Daphnia)-filtration<br>S (Sidae)-filtration<br>Stationary Suspension<br>Raptorial |

### Statistical analyses

To visualize how phytoplankton and zooplankton taxa and traits were distributed on the landscape, we used the percentage of lakes in which they occurred. We tested for differences in taxonomic or functional composition between the three regions using the constrained ordination technique Canonical Analysis of Principal Coordinates (CAP BiodiversityR package; Anderson and Willis 2003). Using the CAP leave-one-out allocation success (% correct, Anderson and Willis 2003) we assessed the distinctiveness of regional composition using the proportion of correct allocation, which can be interpreted as the strength of the compositional differences between regions. Prior to the CAP ordination, taxonomic composition was Hellinger-transformed to reduce the effect of double zeros (Legendre and Gallagher 2001) and Bray-Curtis distance was used for the CAP ordination.

To test whether the distributions of phytoplankton and zooplankton communities were coupled across the landscape, we used a hierarchical framework (Figure 2). First, we tested for significant coupling between the composition of the two groups at taxonomic and functional trait levels using a Procrustes analysis (Mardia et al. 1980) on the subset of lakes (48) for which both phytoplankton and zooplankton were sampled. Specifically, we tested the degree of concordance between the PCA ordinations of phytoplankton and zooplankton taxonomic and functional community compositions. If a significant coupling was observed, we used the residuals from a distance-based redundancy analysis (dbRDA, Legendre and Legendre 1998) to sequentially control for the effect of water quality, morphometry and space (using between-lake distance based on latitude and longitude coordinates) and to test at each step whether significant coupling was observed (Figure 2). For this analysis, species composition was Hellinger transformed (Legendre and Legendre 1998, Legendre and Gallagher 2001), and functional trait composition was logit transformed. We tested the significance of the Procrustes statistic using a permutation procedure (9999 simulation, protest function in vegan R package, Oksanen et al. 2015).

**Figure 2:**
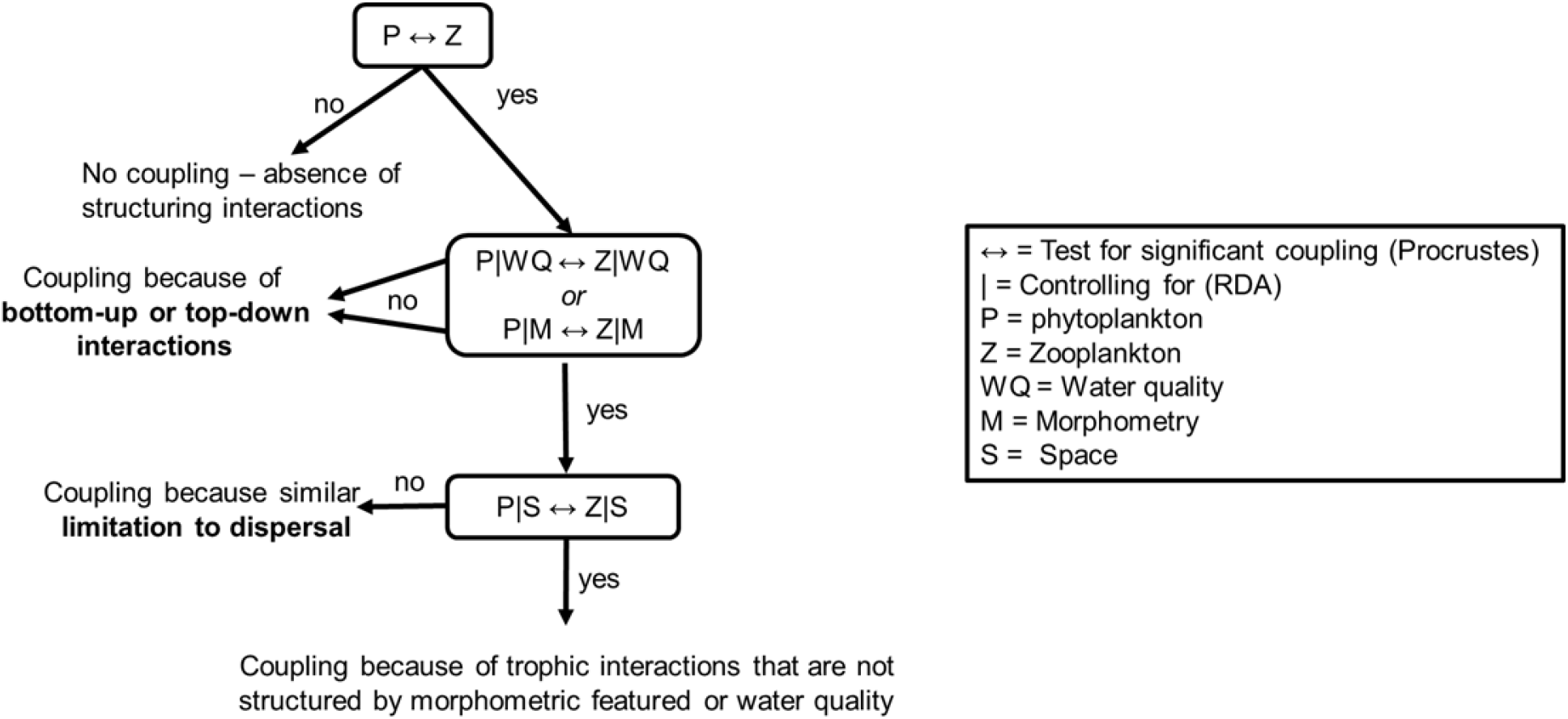
Conceptual framework used to test for a significant coupling between zooplankton and phytoplankton communities using taxonomy and functional traits, while controlling for environmental factors (water quality WQ, morphometry M) and dispersal limitation (space S).

To directly evaluate the relative importance of environmental and spatial variables as drivers of the taxonomic and functional composition of phytoplankton and zooplankton, we used distance-based redundancy analysis for taxonomic composition (dbRDA, Legendre and Legendre 1998) and multiple regression for each functional trait; both followed by variation partitioning (Borcard et al. 1992). We separated the environmental variables into two groups; water quality (WQ with Chl-a, TP, TN, DOC, water color, pH, temperature) and morphometry (M with lake area and depth). Prior to the RDAs and multiple regressions, we used a stepwise selection (based on AIC) of variables within each group of environmental variables. We identified shared variation between the two groups of environmental variables (WQ+M), and between environmental variables and space (S) as: WQ+S and M+S to determine whether environmental variables were spatially structured. Finally, to visualize the relationship between phytoplankton and zooplankton taxa and functional traits, we used an RDA with all taxa and all the functional traits combined.

## Results

### Biogeographical patterns

We observed important environmental differences between the three regions (Figure 1, Table 1). The first principal component (PC) mainly differentiated the Abitibi and Chicoutimi regions from the Schefferville region and represented differences that reflect latitude between the regions: lakes in Abitibi and Chicoutimi being warmer, darker (higher CDOM) with higher concentrations of chlorophyll a, dissolved organic carbon (DOC) and nutrients (TN and TP). The second PC axis differentiated the Chicoutimi and Abitibi regions and was mainly related to morphometric differences: lakes in the Abitibi region were shallower, with flatter catchments and also with higher nutrient concentrations. Lakes in the Schefferville region were spread across the second PC axis indicating that lake morphometry and catchment characteristics were highly variable in this region.

The average percent occurrence (Figure 3a) was 24% for phytoplankton taxa (median 15%) and 21% for zooplankton taxa (Figure 3b, median 8%). Four phytoplankton taxa Mallomonas (90%), Cryptomonas (90%), Dinobryon (82%) and Asterionella (63%), were observed in more than 60% of lakes. For zooplankton Leptodiaptomus (89%), Daphnia (87%), Bosmina (84%) and Holopedium (64%) had occurrences greater than 60%. Of the 56 phytoplankton taxa, 34 (61%) were observed in all three regions, while across all 27 zooplankton taxa, 10 (37%) were observed in all regions. The differences in taxonomic composition between regions were significant for both groups, but were less pronounced for phytoplankton (% correct = 69%, p=0.01, Figure 4a) than for zooplankton (76%, p=0.01, Figure 4b).

**Figure 3:**
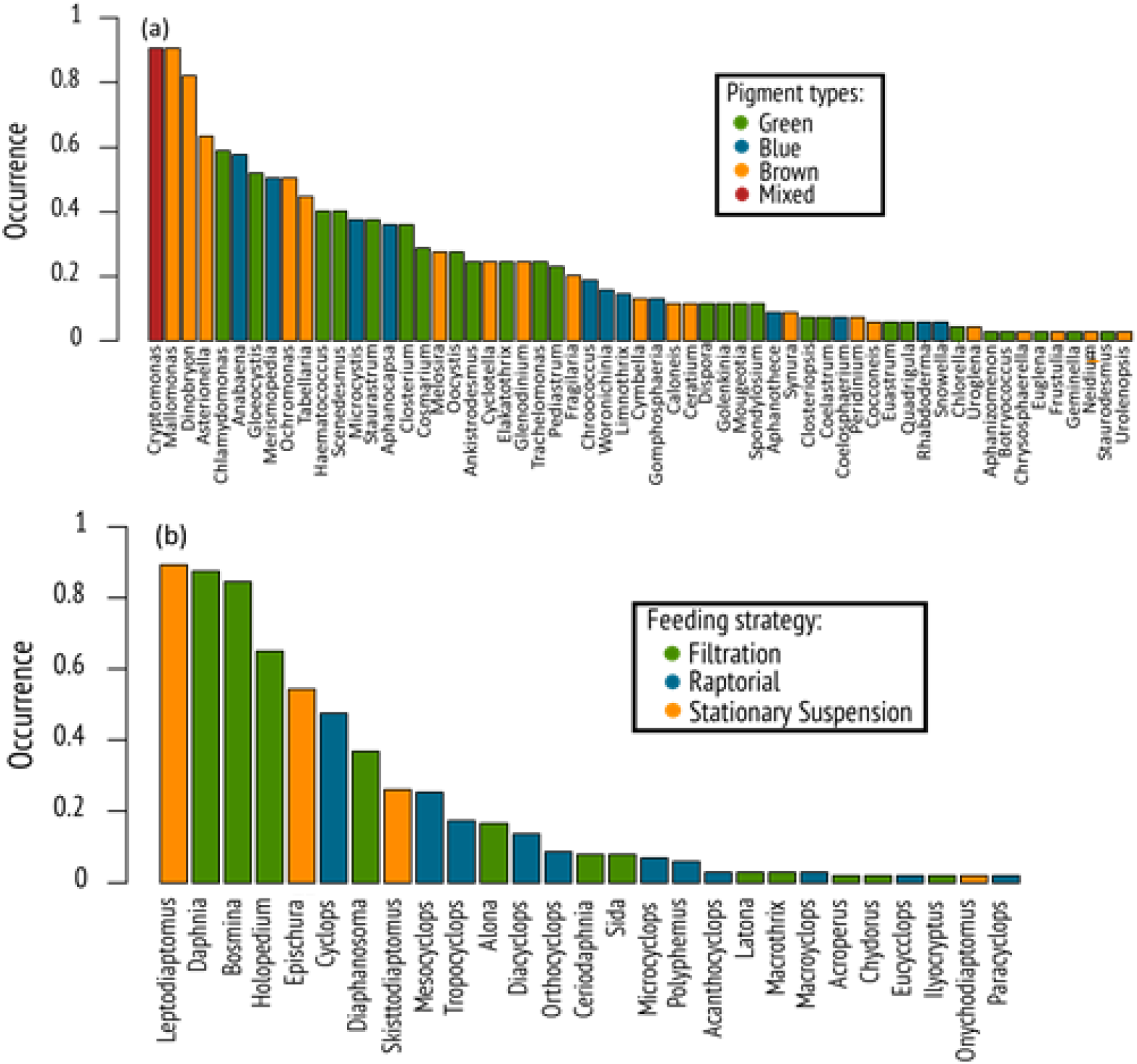
Ranked occurrence (percent of all lakes) of (a) phytoplankton and (b) zooplankton taxa.Bars and dots were coloured by Pigment trait type for phytoplankton and the Feeding strategy trait for zooplankton.

**Figure 4:**
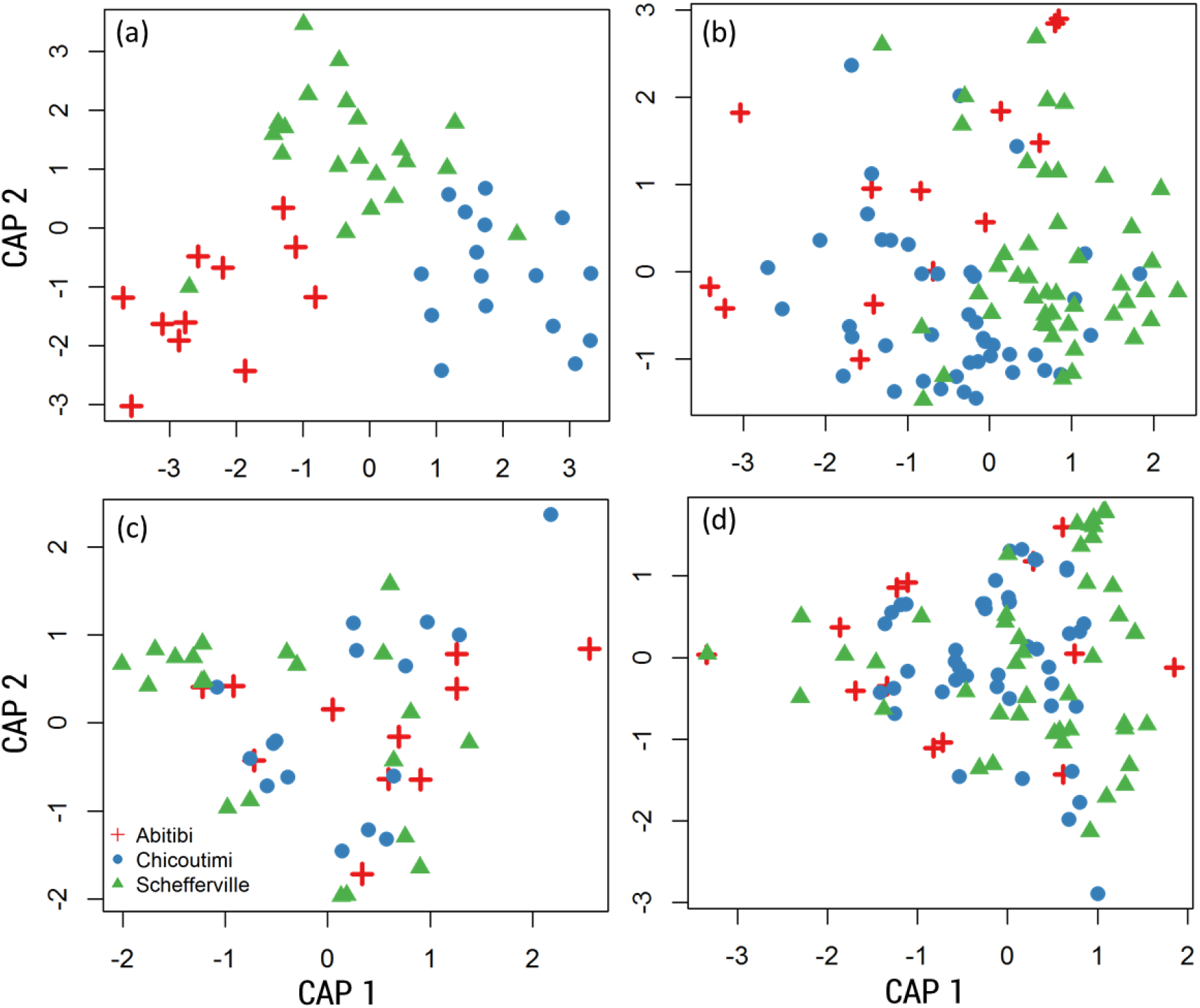
Two-dimensional scatter plot of canonical axes of the CAP ordinations for (a) phytoplankton and (b) zooplankton taxonomic composition and their respective functional trait compositions (c, d). Symbols represent the different regions. Strong regional differences occurred for taxonomic composition of phytoplankton (a, p=0.01) and zooplankton (b, p=0.01), but were not significant for functional trait composition (c and d).

For both phytoplankton and zooplankton, all functional traits were present in all three regions. All phytoplankton functional traits occurred in more than 90% of lakes, with the exception of two traits associated with cyanobacteria: presence of a vacuole for motility (63% of lakes) and the potential to fix nitrogen (58% of lakes). The occurrence of zooplankton functional traits ranged between 18% for carnivores and omnivore-herbivores, and 100% for herbivores, with the average occurrence of zooplankton traits being 64% (median 75%). Functional composition between regions did not differ for either phytoplankton, or zooplankton (% correct = 35%, p=0.69 and 49%, p=0.15 respectively, Figures 3c and 3d).

In the taxonomic Procrustes analyses we found a significant correlation between phytoplankton and zooplankton taxa (Table 3), indicating a taxonomic coupling between the distribution of the two groups. The correlation was not significant after controlling for water quality and morphometry, suggesting that the observed taxonomic coupling could probably be attributed to bottom-up and/or top-down interactions between phytoplankton and zooplankton that are structured by variation in water quality parameters or lake morphometry. On the other hand, using functional traits, there was a significant coupling between the plankton groups only when traits related to trophic interactions between phytoplankton and zooplankton (i.e. without phytoplankton resource acquisition traits; including only motility, colonial and biovolume; Table 2) were used, but not when all traits were considered (Table 3). The significant trait correlation did not remain after controlling for water quality, morphometry or space (independently or together).

**Table 3:** Procrustes rotation analysis of species and the trait dataset correlation coefficients for phytoplankton (Phyto) and zooplankton (Zoo) after controlling (|) for water quality (WQ), morphometry (M) and space (S). Where * = 0.01<p<0.05, ** = 0.001<p<0.01,*** p<0.001, ns = non-significant.

|  | Control for | m12 squared | Correlation |
| --- | --- | --- | --- |
| Taxonomic | Zoo<sub>taxo</sub> ↔ Phyto<sub>taxo</sub> | 0.85 | 0.38*** |
|  | \|wq | 0.86 | 0.37** |
|  | \|m | 0.86 | 0.37*** |
|  | \|s | 0.89 | 0.34** |
|  | \|wq+m+s | 0.92 | ns |
| All traits | Zoo<sub>funct</sub> ↔ Phyto<sub>funct</sub> | 0.92 | ns |
|  | \|wq | 0.92 | ns |
|  | \|m | 0.9 | ns |
|  | \|s | 0.92 | ns |
|  | \|wq+m+s | 0.93 | ns |
| Without phytoplankton resource acquisition traits | Zoo<sub>funct</sub> ↔ Phyto<sub>funct</sub> | 0.92 | 0.28* |
|  | \|wq | 0.95 | ns |
|  | \|m | 0.94 | ns |
|  | \|s | 0.94 | ns |
|  | \|wq+m+s | 0.94 | ns |

### Factors related to community composition

The RDA model explained 4% of the variation in phytoplankton taxonomic composition. For functional composition, up to 27% of variation in the biomass proportion of the different traits was explained in the multiple regressions (Figure 5), but no variation was associated with the distribution of mixotrophy, non-motility, nor biovolume. We observed no direct effect of spatial factors on phytoplankton functional trait variation, but a shared component between water quality and spatial factors indicating that the water quality variables driving the distribution of phytoplankton traits were spatially structured. Water quality variables were most consistently explanatory factors of the phytoplankton functional traits for which variation was explained (blue bars; Figure 5). The exceptions were for the flagellated trait, which responded to lake morphometry, and the mixed pigment trait for which variation was shared between water quality and morphometry. After forward selection, the RDA of phytoplankton taxonomic composition was constrained by TP, CDOM, pH and lake area (Figure 6a). Based on the first axis, differences in taxonomic composition could be mainly explained by lake nutrient status, pH and coloured carbon content (CDOM). Similarly, functional composition also responded to nutrient status and carbon content as well as lake temperature and chlorophyll a concentration according to the first PC axis (Figure 6c). Functional traits related specifically to cyanobacteria (Pigment blue, N fixation and Motility-V) were positively associated with the first axis, while traits (Pigment and mixed brown, Si requiring, Motility-F) related to other key taxonomic groups (including chrysophytes, cryptophytes and diatoms) were negatively associated. In the taxonomic and functional RDA, the first two axes were significant.

**Figure 5:**
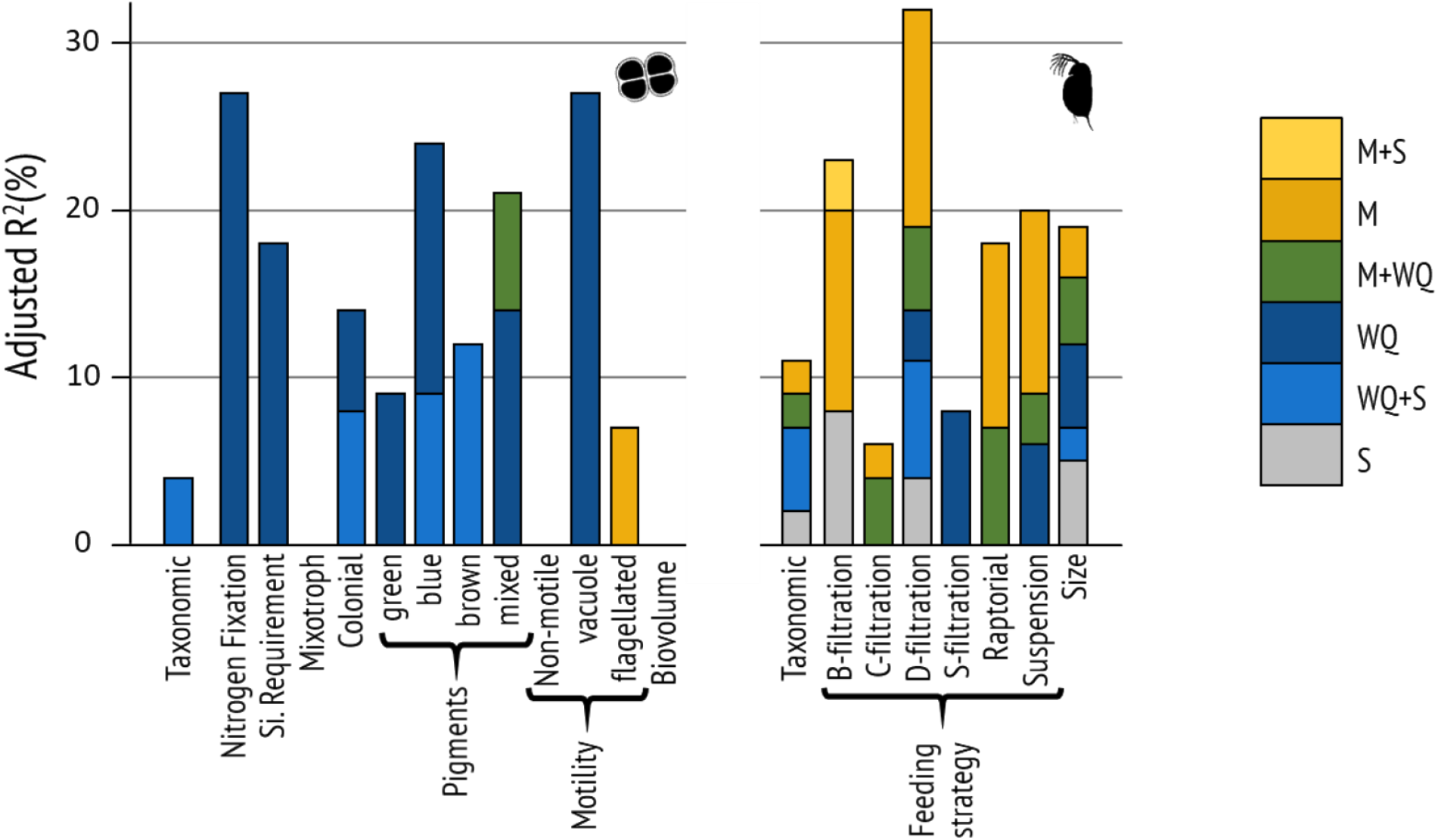
Variation explained (adjusted R^2^) by environmental and spatial factors for phytoplankton (left panel) and zooplankton (right panel) taxonomic composition and each functional trait. Only regression and RDA models that were globally significant are displayed. The variation that was independently explained by a group of variables is represented if significant, and shared variation represents the sum of variation shared between any of the groups of variables, represented with a +. Abbreviations are as follows : morphometry (M) ; water quality (WQ) ; space (S).

**Figure 6:**
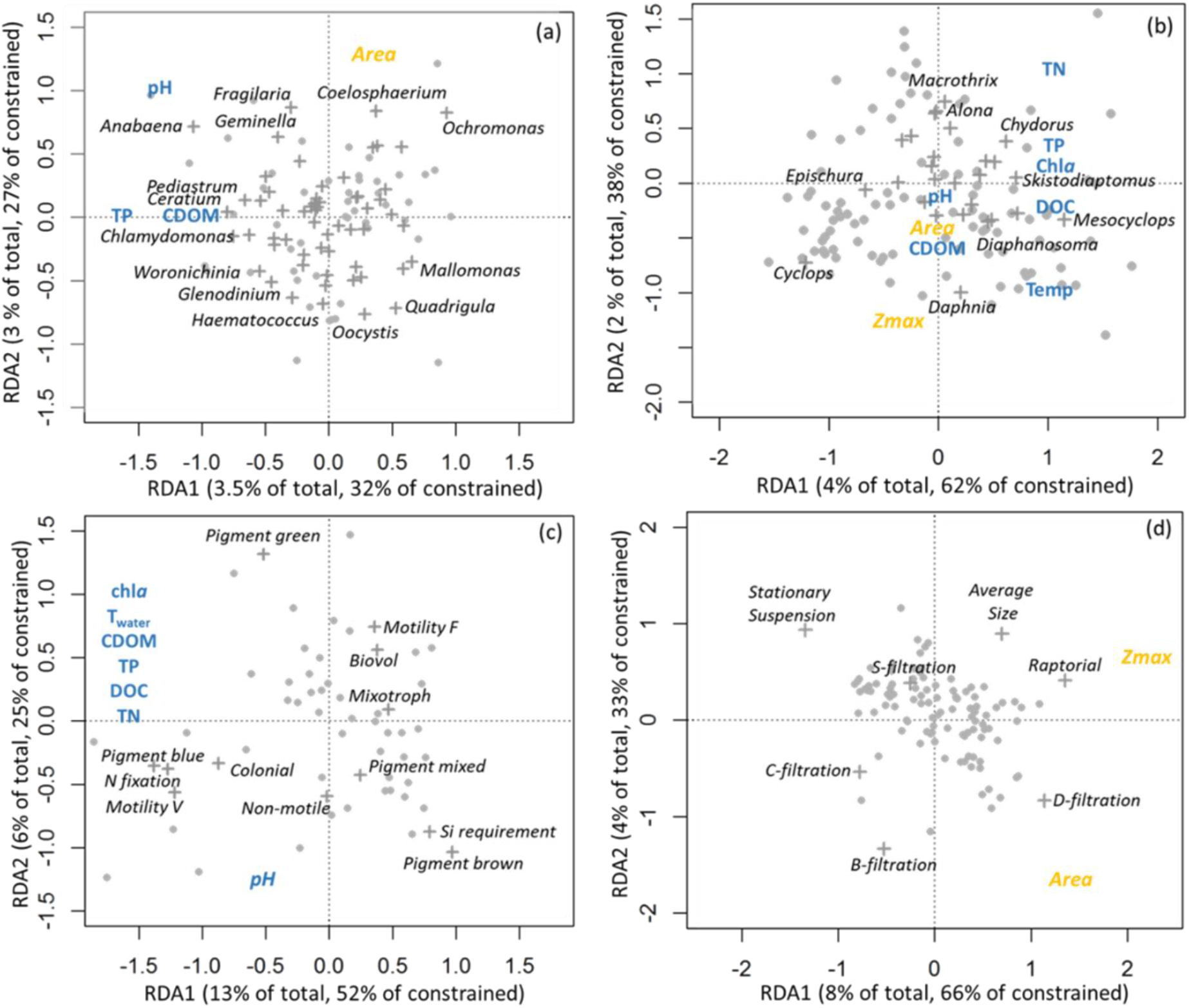
RDA ordination triplots of the (a, c) phytoplankton and (b,d) zooplankton composition classified by (a,b) taxonomy and (c,d) functional (c, d) traits. The RDA was constrained by environmental variables related to water quality (in blue) and by variables related to lake morphometry (in yellow). Taxa and functional trait response variables are represented by crosses and lakes by dots.s. In the taxonomic RDAs (a, b), only taxa with loadings over 0.20 are displayed. Abbreviations are as follows total phosphorus (TP); total nitrogen (TN); dissolved organic carbon (DOC); chlorophyll a (*Chl-a*); coloured dissolved organic matter (CDOM); water temperature (T_water_); lake maximum depth (Zmax); lake area (Area).

For zooplankton, the global RDA models explained 11% of the variation in taxonomic composition (Figure 5), and between 6% and 32% of the variation in the proportion of different functional traits (Figure 5). Similarly, zooplankton variation was shared between water quality and space, but spatial factors also independently explained a significant portion of the variation in zooplankton taxonomic composition, B-filtration, D-filtration and size. For taxonomic composition and most functional traits, a large portion of the variation was attributable to lake morphometry (yellow bars; Figure 5). However, for most traits, and taxonomic composition, some variation was either shared or explained independently by water quality.

In the subsequent RDA of zooplankton taxonomic composition, constrained by morphometric and water quality variables (Figure 6b), variables related to lake productivity (Chla, TP and TN), carbon content (DOC) and temperature loaded on the first axis, while variables relate to lake morphometry (lake depth and area) loaded more strongly on the second axis. For functional traits (Figure 6d), the first axis was mainly related to lake depth, with the second being related to lake area. The raptorial feeding trait was highly correlated with deep lakes and C-filtration with shallow lakes. B-filtration was positively correlated with lake area and size was negatively correlated with lake area while D-filtration was related to large and deep lakes and Stationary suspension to small and shallow lakes. In the taxonomic and functional RDA, the first two axes were significant.

## Discussion

Overall, the distribution of phytoplankton and zooplankton taxa and traits was strongly influenced by water quality and lake morphometry, and to a lesser extent by dispersal limitation across boreal lakes. Although phytoplankton mainly responded to variation in water quality, and zooplankton to lake morphometry, we observed that the distribution of the taxa and traits between the two plankton groups was coupled based on a Procrustes test. Thus, trophic interactions in response to bottom-up and top-down effects in lakes play a significant role in the taxonomic and functional distribution of both phytoplankton and zooplankton. For taxa we also observed that dispersal limitation played a significant role that is consistent with clear taxonomic differences between regions; but these differences did not translate into functional differences between regions.

### Regional differences in composition and environment

Significant differences in phytoplankton and zooplankton taxonomic composition between the three regions indicate a role for environmental factors or dispersal in influencing distributions at the regional scale. On the other hand, functional composition did not differ between regions, indicating that environmental variation or dispersal distances could not preclude a full distribution of possible traits across the entire landscape; instead indicating there is substitution of taxa possessing the same traits (redundancy) in different regions. However, the biogeographical overlap in traits does not mean that functional composition was the same in all lakes. Environmental factors related to water quality and lake morphometry explained a significant amount of variation in relative biomass of most functional traits across both plankton groups (Figure 5), indicating that the control of functional composition acts at sub-regional scales. We also observed that control of plankton taxonomic and functional composition by these lake characteristics was far more important than was the effect of dispersal limitation. Finally, comparing across plankton groups, dispersal limitation was more important for zooplankton than phytoplankton, supporting previous work (Beisner et al. 2006, De Bie et al. 2012), but here now also including functional traits.

### Divergent responses of phytoplankton and zooplankton to their environment

Although both plankton groups responded strongly to environmental factors, the specific variable types accounting for the most variation in taxonomic and functional composition differed between phyto- and zooplankton. Phytoplankton taxonomic and functional trait compositions responded most consistently and strongly to water quality (i.e. their proximal environment). This is to be expected based on previously established strong relationships between composition and lake nutrient status for broad taxonomic groups, groups which were reflected in our functional (pigment) categorization (Watson et al. 1997). On the other hand, zooplankton taxonomic and functional composition both responded more strongly to lake morphometry. However, some water quality effects were also evident for zooplankton, indicating overall an integrated response of zooplankton functional and taxonomic composition to their proximal environment (water quality) and habitat characteristics (lake morphometry).

Zooplankton response to water quality could either occur as a direct effect, or as an indirect bottom-up response to changes in phytoplankton community structure. For the more prevalent response of zooplankton to lake morphometry, this reflects variation in habitat characteristics, such as differences in lake physics (e.g. thermal stratification) or lake depth. We consider it most likely that this morphometry effect arose indirectly, operating via the trophic influence of fish predation, as previously observed (O’Brien et al. 2004). Specifically, lake depth influences fish community composition (Jackson and Harvey 1989), with larger volume lakes tending to have longer food chains (Post et al. 2000), thereby modulating the trophic cascade effect on zooplankton through planktivore fish feeding (Carpenter et al. 1985). The variation in zooplankton taxonomic and functional trait composition related to lake morphometry could thus result from local variation in fish composition (data which we did not have). Further evidence comes from the larger proportions of D-filtration in larger, deeper lakes: in larger lakes the presence of an extra trophic level (total of 4-levels) decreases fish zooplanktivory, resulting in reduced top-down pressure on large herbivorous cladocera, which are preferred prey for fish (Christoffersen et al. 1993). Repercussions throughout the community were observable in the ordination biplots (Figure 6d), with reductions in D-filtration being related to an increased proportion of stationary suspension herbivory, dominated by calanoid copepods. Also, the relative biomass of the C-filtration group was negatively related to lake depth, consistent with the fact that most species within this functional feeding type are littoral species and shallow lakes contain greater proportion of habitats that are littoral.

### Effect of trophic interactions on the distribution of phytoplankton and zooplankton

Using the common currency of functional traits, based on a Procrustes test we observed a coupling between the biogeography of phytoplankton and zooplankton only when phytoplankton traits reflecting trophic interactions (grazer avoidance traits) were included. This confirms that the selection of functional traits needs to be guided by the ecological question being posed (Petchey & Gaston 2006). However, this functional coupling was not significant after controlling for water quality and morphometry, indicating that top-down and bottom-up interactions structured trophic interactions between the two groups leading to a coupled trait distribution of phytoplankton and zooplankton on water quality and morphometry gradients. The fact that dispersal was not a factor in this coupling is not surprising given that all traits were distributed across the landscape with no differences between the three regions. The observed coupling, that we attribute to trophic interactions, supports other functionally based models to describe coupled plankton dynamics, also influenced by water quality and morphometric factors, through time (e.g. PEG model). Our results further suggest that trophic interactions are consistent enough at the landscape level to result in a spatial functional coupling of phytoplankton and zooplankton.

Contrary to our expectations, the coupling observed between phyto- and zooplankton was stronger at the taxonomic level, suggesting that trophic interactions play a larger role for taxonomic than functional composition. This is further supported by the concordant taxonomic response to phosphorus (TP, Figure 6 a and b) indicating that phosphorus enrichment affects phytoplankton taxa composition, which triggers a response in the zooplankton, as would be expected with changes in edibility of phytoplankton along such a nutrient gradient (e.g. Watson et al. 1997). However, the coupling was much more reduced when controlling for space, suggesting that similarities in dispersal limitation is a stronger driver of phytoplankton and zooplankton distributions than is the environment. This would explain why at the scale of this study, we observed a stronger coupling for taxa, as there was no effect of dispersal on trait distribution. It is also important to acknowledge that the coupling between plankton groups that we attribute to trophic interactions structured by environment could also be the result of the common response of both groups to the same gradient, for example a TP gradient. Unfortunately, mechanistic links such as these are difficult to verify with observational studies like ours at these spatial scales, and would require smaller-scale experimental study to fully verify. In any case the importance of trophic interaction between phytoplankton and zooplankton have already been demonstrated (Sommer et al. 1986, 2012).

The goal of including functional traits in our study was to assess whether biogeographical coupling would be more easily observed using a more mechanistic aggregation of organismal differentiation than is done by pure taxonomy. To this end, we selected functional traits that are directly representative of interactions between phytoplankton and zooplankton and in resource acquisition. Counter-intuitively we found that evidence for coupling was stronger at the taxonomic level, compared to the functional. However, our results suggest that part of the explanation resides in the fact that taxa were dispersal-limited, which strengthens the correlation between the adjacent trophic levels, while traits were conserved at the scale of our study because of functional redundancy among taxa. Hence, for the suite of feeding related functional traits used in this study, we found support for direct reciprocal influences of the zooplankton and phytoplankton community playing a role in structuring the distribution of plankton across boreal lakes such that strong trophic connections in individual lakes (Porter 1977, Sterner 1989), which constitutes the basis of the main pathway for matter and energy transfer in aquatic environments seem to constrain their joint biogeographical distributions. This result has implications for the predictability regarding large-scale change in plankton taxonomic or functional compositions across lake landscapes with the spread of invasive species or the northward migration of species with climate warming. Responses of pelagic food webs to anthropogenic changes in ecosystem parameters across landscapes of lakes should be driven by a combination of top-down and bottom-factors for taxonomic composition, but with a relative resilience in functional trait composition.

## Data accessibility

Data are available online: 10.5281/zenodo.5556991

## Supplementary material

Script and codes are available online: https://doi.org/10.6084/m9.figshare.3457043.v4, XXXXlink or DOI of the webpage hosting the script and codes (DOI for ZOO and PHYTO)

## Acknowledgements

We thank Monique Arianne Resende and Marilyne Robidoux for help in the lab and field and Roy Nahas for the GIS work. Funding from the Fonds de recherche du Québec - Nature et Technologies (FRQNT) strategic clusters: Groupe de recherche interuniversitaire en limnologie et en environnement aquatique (GRIL) and the Québec Centre for Biodiversity Science (QCBS) is gratefully acknowledged. BEB and PDG acknowledge funding from the Natural Sciences and Engineering Research Council of Canada (NSERC) in the form of Discovery Grants and a Research Chair in Carbon Biogeochemistry of Boreal Aquatic Systems (CarBBAS) co-funded by Hydro-Québec to PDG. Version 4 of this preprint has been peer-reviewed and recommended by Peer Community In Ecology (https://doi.org/10.24072/pci.ecology.100082)

## Conflict of interest disclosure

The authors of this preprint declare that they have no financial conflict of interest with the content of this article.

## Notes

https://doi.org/10.5281/zenodo.5556991

### Competing Interest Statement

The authors have declared no competing interest.

### Summary of Updates

This version of the manuscript has been revised following review process in PCI

